# YTHDF1 Facilitates Lung Adenocarcinoma Progression via Promotion of EEF1G Translation in a m6A-Dependent Manner

**DOI:** 10.1101/2024.09.13.612607

**Authors:** Lihong Wang, Qihong Sheng, Xiaoyu Wang, Hongjuan Yue, Qian Wang, Mei Zhang, Junling Ma, Ling Wu, Jiaojiao Zhang, Zishuo Cheng, Weifang Yu, Ting Liu, Jia Wang

## Abstract

Lung adenocarcinoma (LUAD) is a malignant tumor with high morbidity and mortality worldwide, and overall survival rates for LUAD patients remain unimproved. RNA modification is a key process in post-transcriptional gene regulation in epigenetics, with N6-methyladenosine (m6A) being a common RNA modification. The molecular mechanisms of LUAD are unclear, but evidence suggests that m6A RNA methylation plays a significant role. This study aimed to clarify the role of YTHDF1 in LUAD development and pathogenesis. These findings confirmed that YTHDF1, a m6A reader protein, is highly expressed in LUAD tissues and is correlated with tumor differentiation and TNM stage. The results of functional loss experiments in LUAD cell lines revealed that downregulating YTHDF1 inhibits proliferation, migration, and invasion and induces apoptosis, with opposite effects observed upon YTHDF1 upregulation. *In vivo*, YTHDF1 knockout suppressed LUAD xenograft growth. RNA-seq, MeRIP-seq, RIP-seq, and bioinformatics analyses identified EEF1G as a downstream target of YTHDF1 in LUAD, and high expression of EEF1G was confirmed. The interaction between YTHDF1 and EEF1G was validated through RIP-qPCR, Co-IP and Co-IF assays. The overexpression of EEF1G in LUAD cells partially counteracts the tumor suppression induced by YTHDF1 silencing, and the knockdown of EEF1G has the opposite effect, further confirming the regulatory relationship. In summary, this study describes a novel YTHDF1/EEF1G regulatory pathway in which YTHDF1 promotes LUAD progression by recognizing and binding to the m6A-modified mRNA of EEF1G, accelerating its translation, suggesting that YTHDF1 may be a potential biomarker and therapeutic target.

## Introduction

Lung cancer (LC) remains the leading cause of cancer-related death^1^. Pathologically, LC can be categorized into two primary subtypes: small cell lung cancer (SCLC) and non-small cell lung cancer (NSCLC), accounting for 15% and 85% of all cases, respectively^2^. NSCLC can be further subdivided into squamous cell carcinoma, adenocarcinoma, and large cell carcinoma^3^. Lung adenocarcinoma (LUAD) has emerged as the most common respiratory tumor, with its incidence among younger individuals increasing annually^4^. Despite significant advancements in the treatment of LUAD, including the use of targeted therapies for late-stage metastatic patients, resulting in long-term survival rates of 5 to 7 years or more, the exploration of potential mechanisms of LUAD and the identification of effective therapeutic targets remain critical challenges in the field.

Epigenetics is generally defined as chemical modification, including DNA and RNA methylation, histone modification, noncoding RNA modification, and chromatin rearrangement^5^. m6A RNA methylation refers to methylation at the 6th nitrogen atom of adenine (A) bases in RNA, which is most abundant in mRNAs and long noncoding RNAs (lncRNAs), thus serving as a major epitranscriptomic regulator^6^. It influences the fate and expression of RNA by modulating RNA splicing, nuclear export, stability, and translation, playing crucial roles in the pathogenesis of various diseases, including cancer^7^. m6A methylation is dynamic and reversible and is regulated by a series of m6A modification enzymes, which can be categorized into three types: writers (methyltransferases), erasers (demethylases), and readers (RNA-binding proteins)^8^. An increasing body of evidence suggests that dysregulation of m6A modification-related proteins is involved in various cancers and plays important roles in tumor cell proliferation, differentiation arrest, survival, carcinogenesis, and metastasis.

m6A readers are crucial m6A-binding proteins, typically containing the YTH domain, with YTH proteins being the most thoroughly studied m6A-binding proteins, including YTH domain family protein 1 (YTHDF1), YTHDF2 in the cytoplasm, and YTHDF3, YTH domain containing 1-2 (YTHDC1-2) in the nucleus^9^. These readers directly recognize m6A sites through their YTH domain and determine the fate of target mRNAs by regulating their splicing, nuclear export, translation, and/or stability^10^. In recent years, numerous studies have shown that YTHDF1 plays a key role in tumorigenesis and metastasis through diverse mechanisms, such as promoting translation or regulating mRNA stability. For example, YTHDF1 can bind to m6A sites surrounding stop codons in mRNAs; recruit translation initiation complexes, including eukaryotic translation initiation factor 3, eukaryotic translation initiation factor 4E, eukaryotic translation initiation factor 4G, poly(A)-binding protein, and the 40S ribosomal subunit; and be recruited to promote the translation of target RNA^11^. Chen et al. reported that YTHDF1 can recognize and bind to FOXM1 mRNA modified by m6A, accelerating the translation of FOXM1 and promoting breast cancer metastasis^12^. Research by Chen et al. also indicated that the m6A reader YTHDF1 promotes gastric carcinogenesis and metastasis via m6A-dependent translation of the USP14 protein^13^. YTHDF1 is highly expressed in cervical cancer and is closely associated with poor prognosis, promoting the translation of RANBP2 in a m6A-dependent manner to increase the growth, migration, and invasion of cervical cancer cells^14^. Several studies have explicitly demonstrated that YTHDF1 can promote high expression in various tumors, including ovarian cancer, bladder cancer, rectal cancer, and hepatocellular carcinoma, and is associated with poor prognosis. In LUAD, although there have been reports that YTHDF1 expression is significantly increased in tumor tissue compared with adjacent normal tissue and is closely associated with poor prognosis, the specific mechanism remains unclear. Moreover, YTHDF1 has attracted extensive attention as a potential therapeutic target. Therefore, this study aimed to investigate the role and mechanism of action of YTHDF1 in LUAD to explore new therapeutic targets for LUAD.

In this study, we analysed data from the TCGA database and the GEO database (GSE43458) and found that YTHDF1 is highly expressed in LUAD tissues, and this finding was further confirmed through the analysis of clinical tissue samples and cell lines. Our clinical data revealed that the level of YTHDF1 expression is positively correlated with the TNM stage of LUAD tumors and negatively correlated with the degree of tumor differentiation. Subsequent functional experiments demonstrated that YTHDF1 promotes the occurrence and development of LUAD cells both *in vitro* and *in vivo*. By integrating RNA-seq, MeRIP-seq, and RIP-seq with bioinformatics analysis techniques, EEF1G was identified as a key m6A modification target of YTHDF1 in LUAD cells. Further experiments demonstrated that YTHDF1 promotes the translation of EEF1G mRNA by recognizing m6A sites on it, increasing its protein expression, and plays an oncogenic role in the pathogenesis of LUAD.

## Materials and methods

### Patient samples

This study collected wax blocks from 52 LUAD patients and their adjacent normal tissues, which were surgically resected at the Department of Thoracic Surgery at the First Hospital of Hebei Medical University in Shijiazhuang, China, between 2017 and 2021. All included patients with LUAD were confirmed by postoperative pathology. None of the patients had received radiotherapy, chemotherapy, or biological treatment before surgery, and all patients signed informed consent forms. The study was approved by the Ethics Committee of the First Hospital of Hebei Medical University (Ethical Approval Number: 20220405). The research conforms to the Declaration of Helsinki.

### Immunohistochemistry (IHC)

Routine paraffin sections from human and nude mouse tissues were dewaxed and hydrated. After antigen retrieval using Tris-EDTA antigen retrieval solution (BOSTER-BIO, AR0023-1, pH=9.0), immunohistochemical analysis was performed via a rabbit/mouse two-step detection kit (Beijing ZSBG-Bio Co., Ltd., Beijing, China) according to the manufacturer’s instructions. Antibodies against YTHDF1 (diluted 1:200; Proteintech, Wuhan, Hubei, P.R.C.; 17479-1-AP), Ki67 (diluted 1:50; Proteintech; 27309-1-AP), and EEF1G (diluted 1:500; Proteintech; 68148-1-Ig) were used for immunohistochemical staining. The results were analysed via a double-blind method; positivity was determined via comprehensive scoring of the staining intensity and range. The staining intensity was assessed as follows: 0 points (pale blue), 1 point (pale yellow), 2 points (brownish yellow), and 3 points (brown). The staining range was evaluated as 0 points (<5%), 1 point (6%-25%), 2 points (26%-50%), 3 points (51%-75%), or 4 points (76%-100%). When the score is multiplied, a total score of > 6 points is considered positive, and a score of 6 points or less is considered negative.

### Cell culture

The cell lines used in this study included the human normal lung epithelial cell line BEAS-2B and the human LUAD cell lines NCL-H1975, NCL-H1299, and A549. All cell lines were purchased from Wuhan Pricella Biotechnology Co., Ltd., and validated by STR analysis. BEAS-2B cells were cultured in DMEM medium (Gibco, Gaithersburg, MD, USA), and the other three LUAD cell lines were cultured in RPMI 1640 medium (Gibco, USA). The culture medium was supplemented with 10% fetal bovine serum (FBS; Gibco, USA) and 1% penicillin‒streptomycin solution (Solarbio Sciences & Technology Co. Ltd., Beijing, China). All the cells were maintained at 37°C in a 5% CO2 cell culture incubator. Routine Mycoplasma testing was performed via PCR. The cells were grown for no more than 25 passages in total for any experiment.

### Plasmids, siRNAs, shRNAs, transfections and stable cell lines

When the cell density reached 80–90%, the cells were cultured in a cell culture flask or dish. The transfection was performed via Lipofectamine 3000 (Invitrogen, Carlsbad, CA, USA). To knock down YTHDF1, H1975 and H1299 cells were transfected with YTHDF1 siRNA (siYTHDF1) or a negative control (siNC) (GenePharma Co., Ltd., Shanghai, China). For YTHDF1 overexpression, H1975 and A549 cells were transfected with the pcDNA3.1-YTHDF1 plasmid (YTHDF1) or negative control (Vector) (GenePharma). A549 cells were transfected with EEF1G siRNA (siEEF1G) or negative control (siNC) (GenePharma), and H1975 cells were transfected with the EEF1G overexpression plasmid (EEF1G) or negative control (vector) (GenePharma). The culture medium was replaced every 6–8 hours after transfection. Following transfection, the cells were used for protein/RNA extraction, cell functional assays, or immunofluorescence staining. H1975 cells at a density of 30×10^4^ were incubated with 2 ml of the supernatant for 48 hours and then transfected with YTHDF1 shRNA (shYTHDF1) or a negative control (shNC) (GenePharma). The cells were then selected with 1 μg/ml puromycin, and after 3 days, the selection was continued with 1 μg/ml puromycin until a stably transfected cell line was obtained, which was used for subsequent animal experiments and preparation of sequencing samples.

### RNA isolation and qRT‒PCR

Total RNA was extracted from normal lung epithelial cells and LUAD cells via the RNA-Easy Isolation Reagent (Vazyme Biotech Co., Ltd., Nanjing, China) according to the manufacturer’s instructions. The RNA was reverse-transcribed into cDNA via a PrimeScript RT reagent kit (Takara, Beijing, China). Quantitative real-time PCR (qRT‒PCR) was performed via AceQ Universal SYBR qPCR Master Mix (Vazyme) according to the manufacturer’s instructions, with all samples normalized to β-actin. The procedure was set as follows: predenaturation at 95°C for 5 min and 95°C for 10 s for 45 cycles. The primers used for qRT‒PCR are listed in Supplementary Table 1.

### Western blot and antibodies

Total protein was extracted from the cells via RIPA lysis buffer (Solarbio) supplemented with the protease inhibitor PMSF (Solarbio) at a ratio of 100:1, according to the manufacturer’s instructions. The protein concentration was quantified via a bicinchoninic acid (BCA, Solarbio) protein assay kit. Protein samples of 25–30 μg were loaded onto a 10% SDS‒PAGE gel (Bio-Rad Laboratories Inc., Hercules, CA, USA) and electrophoresed at 80 V for 2 hours. The proteins were then transferred onto a PVDF membrane at a current of (length×width×2) mA for 40 minutes. After being blocked with 5% nonfat milk for 2 hours, the membrane was incubated with the primary antibody overnight at 4°C. The antibodies used for Western blotting were as follows: YTHDF1 (diluted 1:200, Proteintech, 17479-1-AP), EEF1G (diluted 1:1000, Proteintech, 68148-1-Ig), β-actin (diluted 1:1500, ZSBG-Bio, TA-09), and GAPDH (diluted 1:500, Goodhere-Bio, AB-P-R 001). The PVDF membrane was then incubated with the fluorescent secondary antibodies goat anti-rabbit and goat anti-mouse. The immunoreactive protein bands were detected with an Odyssey Scanning System (LICOR Biosciences, Lincoln, NE, USA).

### Cell proliferation assay

For the CCK-8 assay, after the LUAD cell lines were transfected for 48 hours, the cells from each group were digested with trypsin and resuspended, and the cell concentration was adjusted to 3×10^4^/ml. Then, 100 μL of the cell suspension (3000 cells per well) was added to each well, with 4 replicate wells for each group. The wells were labelled for 24 h, 48 h, 72 h, and 96 h and placed in an incubator. At each time point (24 h, 48 h, 72 h, and 96 h), 10 μL of Cell Counting Kit-8 (CCK-8; Dojindo, Tokyo, Japan) reagent was added to the corresponding time points and incubated in the cell incubator for 2 hours. After incubation, the optical density (OD) values were measured at a wavelength of 450 nm via a Promega GloMax luminescence detector (Promega, Madison, WI, USA). The experimental data were recorded, the cell growth curves were plotted, and the experiment was repeated three times to validate the results.

For the colony formation assay, the cells from each group were seeded into 6-well plates at a density of 500 cells per well. The culture medium was changed every 3 days. Following a cultivation period of 7–14 days, the cells were rinsed twice with PBS, fixed with 4% paraformaldehyde for half an hour, and subsequently stained with 0.1% crystal violet for 20 minutes.

### Cell migration assay

For the wound healing assay, when the cell density in each group reached approximately 80–90%, a sterile 200 μL pipette tip was used to gently create scratches on the cell layer. After scratching, the cells were gently washed three times with 2 mL of PBS, and the medium was replaced with FBS-free medium. The scratches were photographed, and the distance at the center of the scratch was measured before the plate was placed back into the incubator. The cells were cultured for an additional 48 hours after scratching and then gently washed twice with 2 mL of PBS. The scratches were photographed again under a microscope, and the distance at the center of the scratch was measured to calculate the wound healing rate.

For the Transwell migration assay, after transfection for 48 hours, the cells from each group were digested with trypsin and resuspended in FBS-free medium, and the cell concentration was adjusted to 4×10^5^/ml. In a 24-well plate, 600 μl of medium containing 10% FBS was added to the lower chamber, and 100 μl of the cell suspension was placed into the upper chamber (Corning Incorporated, Corning, NY, USA), with 4×10^4 cells per insert. The plate was then placed in an incubator at 37°C for 48 hours. After incubation, the Transwell chambers were removed, the culture medium in the wells was discarded, and the chambers were washed three times with PBS. The cells were fixed with 4% paraformaldehyde for 30 minutes and stained with 0.1% crystal violet for 20 minutes. Excess crystal violet was removed by washing with PBS, and five random fields were photographed. The stained cells were counted via ImageJ software (National Institutes of Health, Bethesda, MD, USA).

### Cell invasion assay

For the Transwell invasion assay, the Transwell invasion chambers were precoated with FBS-free medium in the upper chamber before cell seeding and then placed in the cell culture incubator at 37°C for 1 hour of incubation. The remaining steps were the same as those described above for the Transwell migration assay.

### Cell apoptosis assay

For the cell apoptosis assay, 48 hours after transfection, the culture medium was not replaced, and the supernatant and adherent cells from each group were collected and washed twice with PBS. The Annexin V-FITC/PI apoptosis assay kit (NeoBioscience, Shenzhen, China) was used to stain the cells in each group according to the manufacturer’s instructions. After 15 min, the degree of cell apoptosis was analysed via flow cytometry (BD Biosciences, San Jose, CA, USA). Finally, the proportion of apoptotic cells was recalculated via FlowJo software.

### Subcutaneous xenograft mouse models

Ten 4-week-old male BALB/c-nu nude mice (purchased from Beijing Huafukang Biotechnology Co., Ltd.) were randomly divided into a control group (n = 6) and an experimental group (n = 6). H1975 cells stably transfected with shNC or shYTHDF1 (0.5×10^7^ per mouse) were subcutaneously injected into the left thigh skin of the nude mice. Starting from the third day after cell injection, the length and width of the tumors were measured, and the volume was calculated (length × width^2/2). Measurements were taken every two days thereafter. On the 21st day (the tumor volume approached 1000 mm3), the nude mice were euthanized humanely, the tumor masses were excised, and the dissected tumors were weighed and photographed. The data related to the volume and weight of the nude mice were analysed. The tumor tissues were processed for histological examination via paraffin-embedded sections. The above experiments were approved by the Ethics Committee of the First Hospital of Hebei Medical University and were carried out in accordance with the experimental animal care and use system.

### RNA sequencing (RNA-seq)

Total RNA was extracted from H1975 LUAD cells successfully transfected with shYTHDF1 or shNC, and the concentration and quality of the RNA were assessed. The RNA samples were sent to BGI Genomics (Shenzhen BGI Genomics Co., Ltd.) for sequencing. Specifically, the NEBNext® Ultra™ RNA Library Prep Kit was used to construct the library from the qualified RNA and to perform quantitative analysis to validate the library quality. Sequencing was carried out via the Illumina sequencing platform to obtain raw data. Sequence information annotation was completed through data quality control and sequence alignment. Differential expression analysis was performed via the R language DESeq2 and edgeR programs. Genes with log2|fold change| > 1.0 and P < 0.05 were selected.

### Methylated RNA immunoprecipitation sequencing (MeRIP-seq)

Total RNA was extracted from H1975 LUAD cells successfully transfected with shYTHDF1 or shNC, and the resulting RNA library was constructed and sequenced to Novogene. In simple terms, this process involves fragmenting mRNA and using antibodies that specifically recognize the m6A modification to immunoprecipitate RNA fragments with m6A modifications from within the cells. The immunoprecipitated RNA fragments are then subjected to high-throughput sequencing. Combined with bioinformatics analysis, this approach allows for a systematic study of m6A modifications across the entire genome, including statistics on the distribution of differential peaks, motif identification, annotation of differential peaks, and analyses such as GO function enrichment and KEGG pathway enrichment.

### RNA immunoprecipitation and high-throughput sequencing (RIP-seq)

Cells from H1975 cells transfected with shYTHDF1 or shNC were collected 24 hours posttransfection. Following the instructions of the RNA Immunoprecipitation (RIP) Kit (BersinBio, Guangzhou, China; Bes5101), our team used an antibody targeting the protein YTHDF1 (10 μg, Proteintech) and an IgG (10 μg, Proteintech) antibody to precipitate the corresponding RNA‒protein complexes. The captured RNA was then isolated and purified, and the expression of the target genes was detected via qRT‒PCR. Novogene performed ribosomal depletion, library construction, and high-throughput sequencing analysis on the captured RNA. Specifically, RNA fragment distribution and concentration were measured via an Agilent 2100 bioanalyzer (Agilent) and a simpliNano spectrophotometer (GE Healthcare). The NEBNext® Ultra™ RNA Library Prep Kit was used to construct the library from the qualified RNA and to perform quantitative analysis to validate the library quality. The library was then sequenced via the Illumina NovaSeq platform. Filtered raw data were compared to the reference genome, and combined with bioinformatics analysis, a systematic study was conducted on the distribution and annotation of differential peaks across the genome, m6A motif identification, GO function enrichment, and KEGG pathway enrichment analysis.

### Bioinformatics analysis

The mRNA expression of YTHDF1 in LUAD and adjacent normal tissues was analysed via the TCGA (https://www.aclbi.com/static/index.html#/tcga) and GEO (https://www.aclbi.com/static/index.html#/geo) databases. Candidate genes that were initially screened were reanalyzed for GO function enrichment and KEGG pathway enrichment via microBioinformatics-Online Bioinformatics Analysis and Visualization Cloud Platform (http://www.bioinformatics.com.cn/). PrimerBank (https://pga.mgh.harvard.edu/primerbank/) and the Primer Design Tool (https://www.ncbi.nlm.nih.gov/tools/primer-blast/) were used for primer prediction. The overlap of genes selected by RNA-seq, MeRIP-seq, and RIP-seq was analysed via BioVenn (https://www.biovenn.nl/index.php). Gene expression in LUAD and adjacent normal tissues was analysed via GEPIA2 (http://gepia2.cancer-pku.cn/#index). The subcellular localization of genes was investigated via GeneCards (https://www.genecards.org/). The relationships between genes and the survival rate of LUAD patients were analysed via the online Kaplan‒Meier plotter (https://kmplot.com/analysis/index.php?p=service). The m6A methylation levels and YTHDF1 binding enrichment were visualized via IGV software. catRAPID (http://service.tartaglialab.com/page/catrapid_group) was used to predict the binding site motifs between the YTHDF1 protein and EEF1G mRNA.

### Co-Immunofluorescent (Co-IF) assay

For the Co-IF assay, a 10 mm circular coverslip was placed in the center of a six-well plate, followed by the inoculation of H1975 wild-type cells to achieve a cell density of 20–30%. After being cultured for 24 hours, the six-well plate was washed three times with PBS, after which the cells were fixed with 4% paraformaldehyde. The cells were subsequently permeabilized with 0.2% Triton X-100 (Solarbio, T8200) and blocked with 2% bovine serum albumin (BSA-V; Solarbio, A8020). The circular coverslip was then removed and incubated overnight at 4°C with a YTHDF1 antibody (diluted 1:50, Proteintech, 17479-1-AP) and an EEF1G antibody (diluted 1:50, Proteintech, 68148-1-Ig). After the cells were washed three times with PBS, they were incubated for 1 hour at room temperature in the dark with fluorescent secondary antibodies (diluted 1:500; Cy3-conjugated anti-rabbit IgG or FITC-conjugated anti-mouse IgG; Beyotime Biotechnology Co., Shanghai, China). Finally, the cell nuclei were stained with 4’,6’-diamino-2-phenylindole dihydrochloride (DAPI; Beyotime Biotechnology Co.), observed and photographed under an upright fluorescence microscope.

### Co-immunoprecipitation (Co-IP) assay

For the Co-IP assay, H1975 cells were seeded in two 10 cm culture dishes. Once the cells reached confluence, they were collected as pellets. Following the manufacturer’s instructions, the cells were lysed via the Pierce™ Classic Magnetic IP/Co-IP Kit (Thermo Scientific, Waltham, USA; 88804). Subsequently, the YTHDF1 antibody (10 μg, Proteintech) and IgG antibody (10 μg, Proteintech) were added to the lysate separately, and the mixture was rotated and incubated at room temperature for 2 hours. After that, balanced A/G magnetic beads are added, and the mixture is rotated and incubated at room temperature for an additional hour to allow the magnetic beads to adsorb the antibody‒antigen complexes. After the magnetic beads were separated via a magnetic stand, the proteins were eluted, and the eluate was neutralized to a pH of 7. Finally, 5× loading buffer was added for subsequent Western blot analysis.

### Statistical analysis

Data with a normal distribution are presented as the mean ± standard deviation, and data with a nonnormal distribution are presented as the median and interquartile range. Statistical calculations were performed via GraphPad Prism 9.5 (GraphPad Software, La Jolla, CA, USA) and SPSS 25.0 (IBM, Armonk, NY, USA). *P* values were calculated via the Mann‒Whitney U test, t test, Fisher’s exact test or the χ² test, as appropriate. Statistical significance was considered at \**P* < 0.05, \*\**P* < 0.01, and \*\*\**P* < 0.001. All experiments were repeated independently three times.

## RESULTS

### YTHDF1 is upregulated in LUAD samples

To investigate the role of m6A modification-related genes in driving the tumorigenesis of LUAD, we analysed the expression of 21 m6A-related genes in LUAD tissues and adjacent tissues in the TCGA database **(Figure A)**. Four m6A reader proteins (HNRNPA2B1, HNRNPC, YTHDF1, IGF2BP3, and YTHDF2) were significantly more highly expressed in LUAD than in adjacent tissues (*P*<0.001). We further analysed the mRNA expression level of YTHDF1 in LUAD tissues and adjacent tissues in the TCGA database **(Figure B)** and the GEO dataset (GSE43458) **(Figure C)**, which revealed that YTHDF1 mRNA expression was upregulated in LUAD tissues compared with adjacent tissues (*P*<0.01, *P*<0.001). On the basis of the literature, the role of YTHDF1 in LUAD has been less well studied, so YTHDF1 was chosen as the target gene for this research. To further clarify the expression and cellular localization of YTHDF1 in LUAD tissues, we used IHC to detect the expression level of YTHDF1 in 52 pairs of LUAD tissues and adjacent tissues. The results revealed that YTHDF1 was distributed mainly in the cytoplasm, and its expression level in LUAD tissues was significantly greater than that in adjacent tissues (χ²=41.230, *P*<0.001) **(Figure D, Table 1)**. To validate the clinical significance of YTHDF1 in LUAD, we further analysed the correlation between YTHDF1 expression levels and the clinicopathological data of LUAD patients. The results revealed a positive correlation between YTHDF1 expression levels and the TNM stage of LUAD tumors (*P*<0.001) and a negative correlation with the degree of tumor differentiation (*P*<0.05), whereas there was no correlation with patient sex, age, tumor size, or lymph node metastasis (all *P*>0.05) **(Table 2)**. In summary, YTHDF1 is highly expressed in LUAD tissues and is associated with poor patient prognosis.

**Figure 1.**
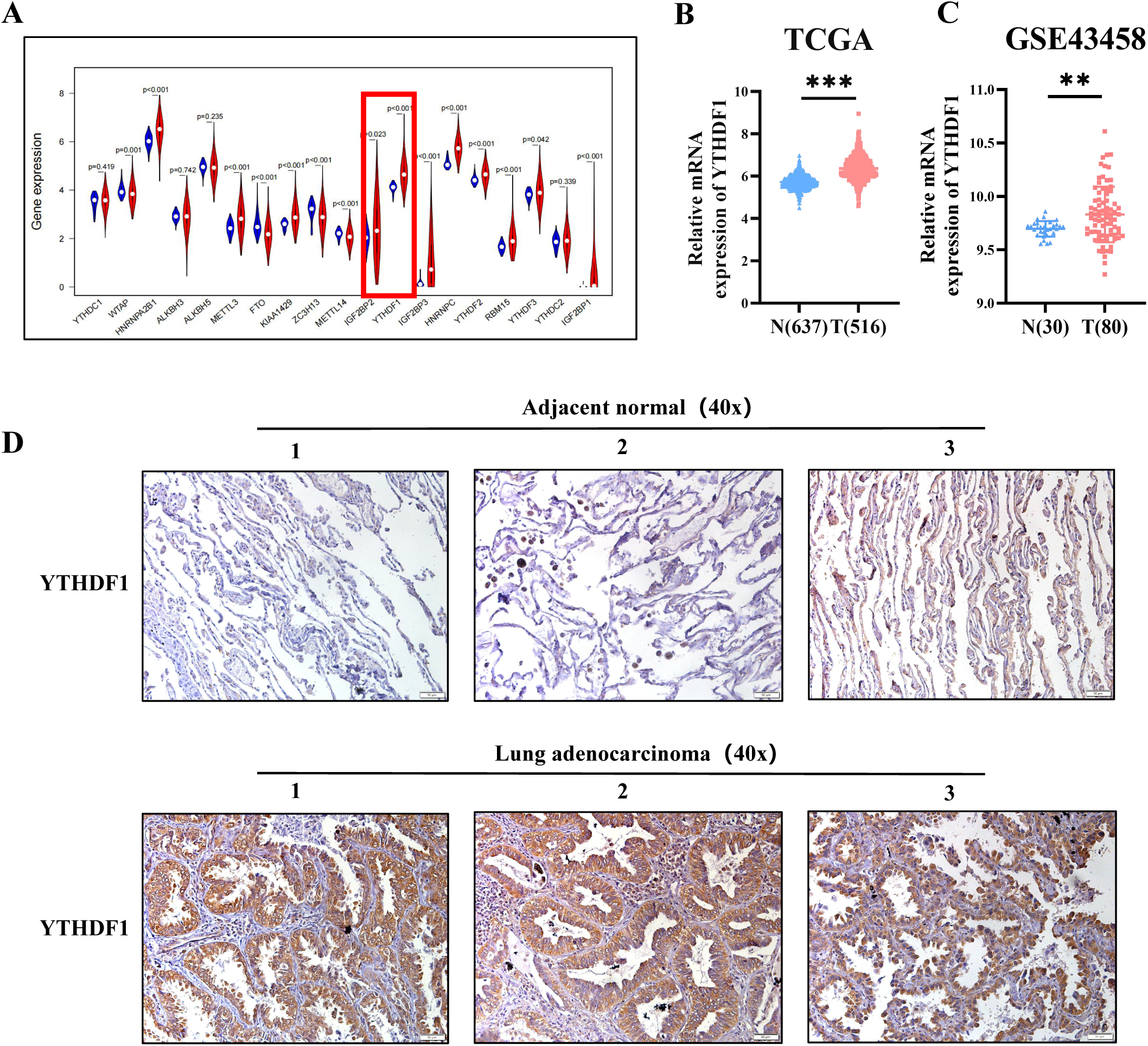
YTHDF1 is upregulated in LUAD samples. **(A)** Expression of 21 m6A-related genes in the TCGA database. **(B, C)** mRNA levels of YTHDF1 according to the TCGA and GSE43458 datasets. **(D)** IHC staining of YTHDF1 in 52 pairs of LUAD tissues and adjacent normal tissues at 40× magnification. Scale bar, 50 μm. The data are shown as the means ± SDs. \*\**P* <0.01, \*\*\**P* <0.001.

**Table 1.**
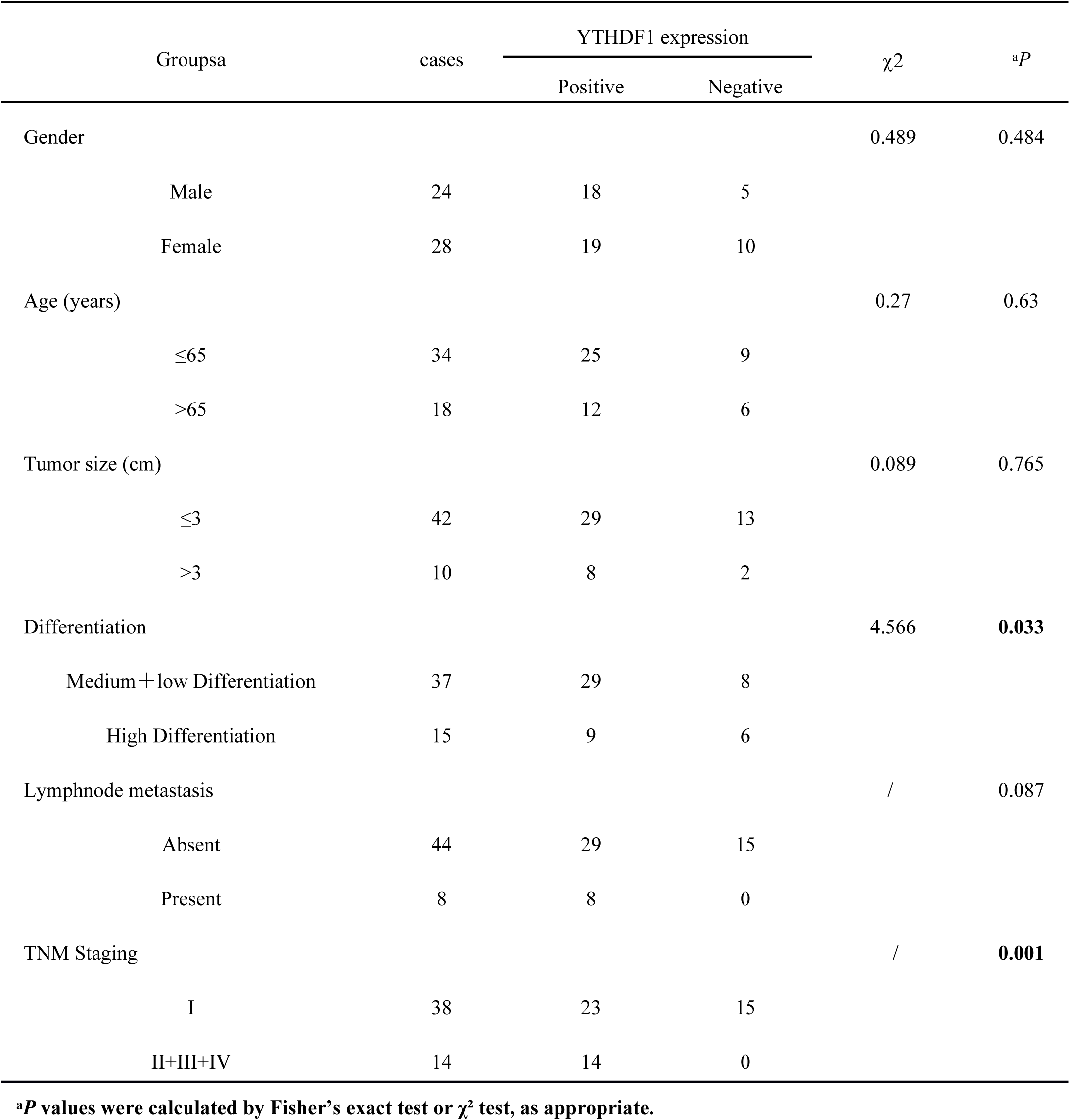
The expression levels of YTHDF1 protein in lung adenocarcinoma and adjacent normal tissues.

**Table 2.**
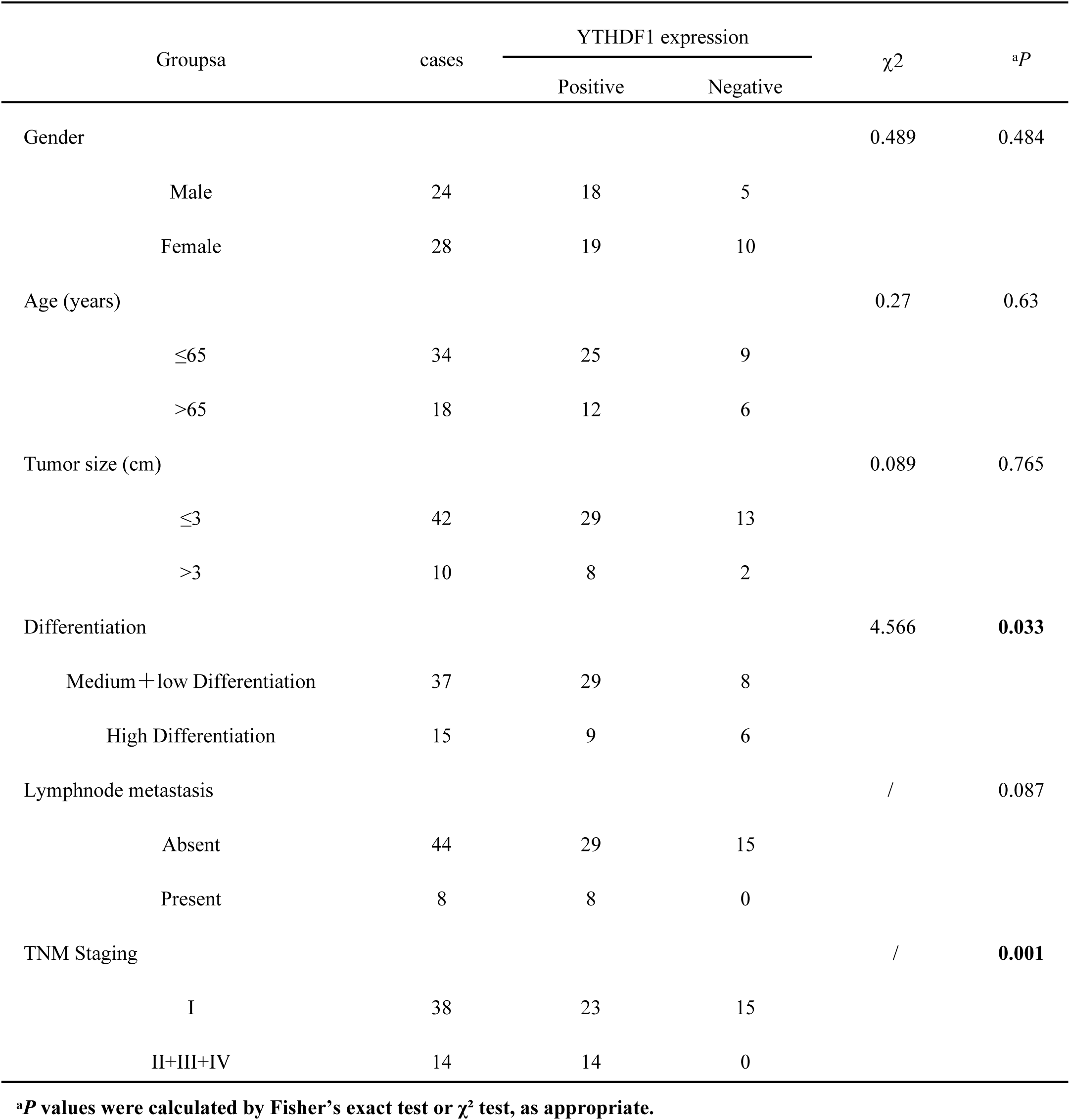
The relationship between the expression of YTHDF1 in lung adenocarcinoma tissues and the clinicoapathological data of patients.

### Knockdown of YTHDF1 suppresses the proliferation, migration, and invasion and promotes the apoptosis of LUAD cells *in vitro*

We transduced siRNAs targeting YTHDF1 into the LUAD cell lines H1975 and H1299 to knock down the expression of YTHDF1. After 48 h of transduction, we detected the mRNA and protein expression levels of YTHDF1 to verify the knockout efficiency via qRT‒PCR and Western blotting, and the results revealed that the siRNA significantly suppressed the expression of YTHDF1 in H1975 and H1299 cells **(Figure 2A‒D)**. CCK-8 assays demonstrated that knocking down YTHDF1 substantially inhibited the proliferation (all *P*<0.001) of the LUAD cell lines H1975 and H1299 *in vitro* **(Figure 2E, F)**. In addition, wound-healing and transwell migration assays revealed that the migration of LUAD cells was significantly suppressed upon YTHDF1 depletion *in vitro* (all *P*<0.01) **(Figure 2G, H, K, L)**. Next, Transwell invasion assays were used to assess whether YTHDF1 affects the invasion of LUAD cells, and the results indicated that YTHDF1 knockdown markedly reduced the invasion of H1975 and H1299 cells *in vitro* (all *P*<0.001) **(Figure 2M, N)**. We also found that YTHDF1 knockdown induced the apoptosis of LUAD cells *in vitro* (all *P*<0.01) **(Figure 2I, J)**. These results suggested that the knockdown of YTHDF1 inhibited LUAD cell proliferation, migration, and invasion but promoted apoptosis.

**Figure 2.**
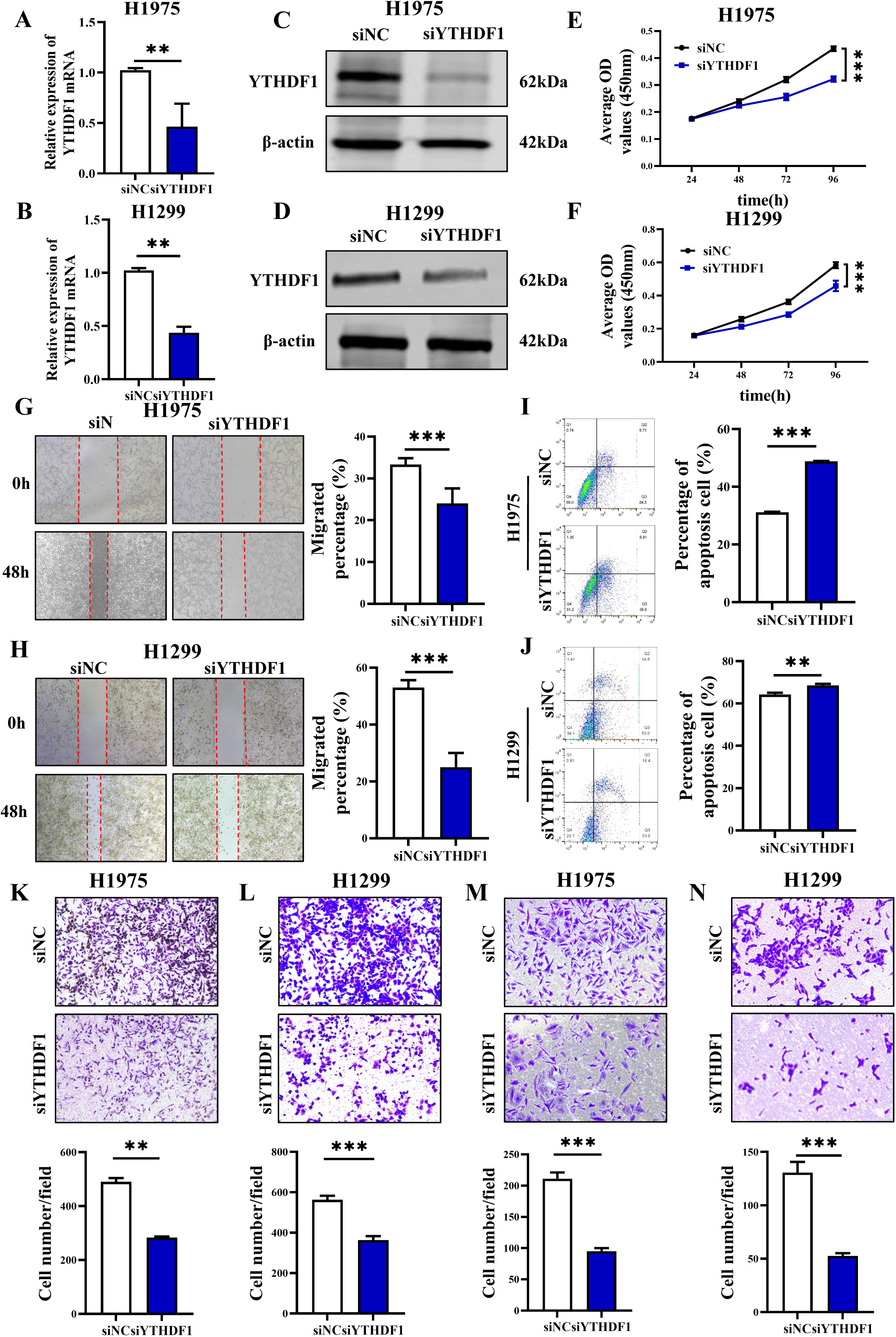
Knockdown of YTHDF1 suppresses proliferation, migration, and invasion and promotes the apoptosis of LUAD cells *in vitro*. **(A, B)** Relative YTHDF1 mRNA expression levels in H1975 and H1299 cells after transduction with scramble siNC or siYTHDF1. **(C, D)** Relative YTHDF1 protein expression levels in H1975 and H1299 cells after transduction with scramble siNC or siYTHDF1. **(E, F)** CCK-8 assay of H1975 and H1299 cells transfected with siNC or siYTHDF1. **(G, H)** Wound-healing assay of H1975 and H1299 cells transfected with siNC or siYTHDF1. Scale bar, 200 μm. **(K, L)** Transwell migration assay of H1975 and H1299 cells transfected with siNC or siYTHDF1. Scale bar, 100 μm. **(M, N)** Transwell invasion assay of H1975 and H1299 cells transfected with siNC or siYTHDF1. Scale bar, 100 μm. **(I, J)** The apoptosis rates of H1975 and H1299 cells transfected with siNC or siYTHDF1 were determined by flow cytometry. The data are shown as the means ± SDs. \**P* <0.05, \*\**P* <0.01, \*\*\**P* <0.001.

### Overexpression of YTHDF1 promotes the proliferation, migration, and invasion and suppresses the apoptosis of LUAD cells *in vitro*

To further explore the biological role of YTHDF1 in LUAD, pcDNA3.1-YTHDF1 (YTHDF1) and pcDNA3.1-vector (Vector) were transfected into the human LUAD cell lines H1975 and A549. First, we verified the efficiency of YTHDF1 overexpression in LUAD cell lines. Through Western blot and qRT‒PCR, the protein and mRNA levels of YTHDF1 were shown to be increased **(Figure 3A-D)**. The effects of YTHDF1 on cell proliferation, migration, invasion and apoptosis were subsequently examined. The results of the CCK-8 assays revealed that, compared with the negative control (vector), the overexpression of YTHDF1 promoted the growth of LUAD cells *in vitro* (all *P*<0.001) **(Figure 3E, F)**. Next, we used wound-healing and Transwell migration assays to evaluate whether YTHDF1 can affect the migration of lung cancer cells and found that the upregulation of YTHDF1 significantly promoted the migration ability of H1975 and A549 cells *in vitro* (all *P*<0.01) **(Figure 3G, H, K, L)**. The potential impact of YTHDF1 on LUAD cell invasion was then investigated via transwell invasion assays. The results revealed that YTHDF1 upregulation significantly promoted H1975 and A549 cell invasion *in vitro* (all *P*<0.001) **(Figure 3M, N)**. We also discovered that the overexpression of YTHDF1 inhibited the apoptosis rate of LUAD cells *in vitro* (all *P*<0.01) **(Figure 3I, J)**. These results suggested that the overexpression of YTHDF1 promoted LUAD cell proliferation, migration, and invasion, whereas it inhibited apoptosis.

**Figure 3.**
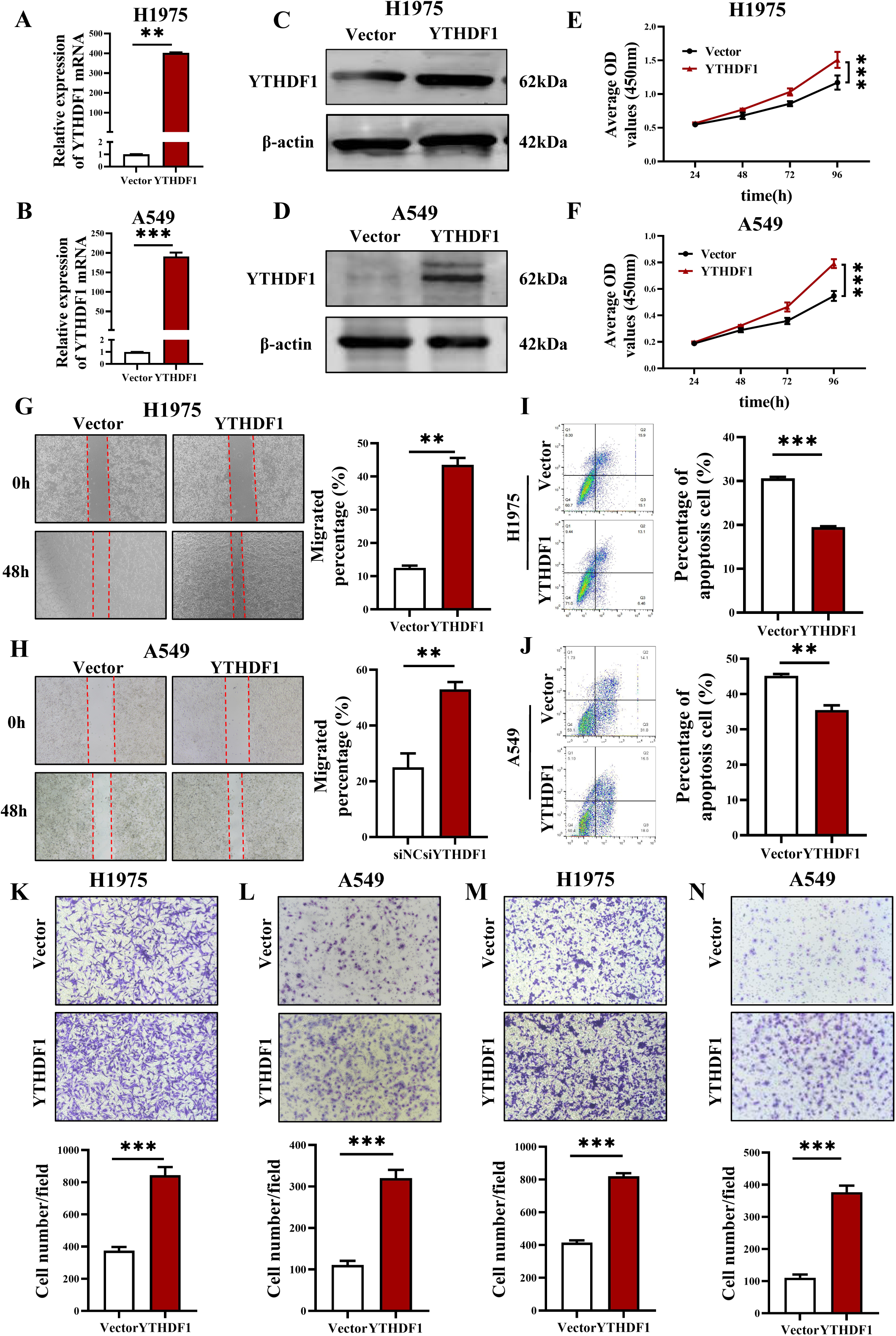
The overexpression of YTHDF1 promotes the proliferation, migration, and invasion and suppresses the apoptosis of LUAD cells *in vitro*. **(A, B)** Relative YTHDF1 mRNA expression levels after the transduction of scramble vector and YTHDF1 in H1975 and H1299 cells. **(C, D)** Relative YTHDF1 protein expression levels after the transduction of scramble vector and YTHDF1 in H1975 and H1299 cells. **(E, F)** CCK-8 assay of H1975 and H1299 cells transfected with vector and YTHDF1. **(G, H)** Wound-healing assay of H1975 and H1299 cells transfected with vector and YTHDF1. Scale bar, 200 μm. **(K, L)** Transwell migration assay of H1975 and H1299 cells transfected with vector and YTHDF1. Scale bar, 100 μm. **(M, N)** Transwell invasion assay of H1975 and H1299 cells transfected with vector and YTHDF1. Scale bar, 100 μm. **(I, J)** The apoptosis rates of H1975 and H1299 cells transfected with vector and YTHDF1 were determined by flow cytometry. The data are shown as the means ± SDs. \**P* <0.05, \*\**P* <0.01, \*\*\**P* <0.001.

### YTHDF1 regulates the growth of LUAD *in vivo*

We created a tumor-bearing nude mouse model with low expression of YTHDF1 and a corresponding negative control to investigate the role of YTHDF1 in LUAD *in vivo*. After successfully transfecting the H1975 cell lines with shRNA-YTHDF1 (shYTHDF1) or shRNA-negative control (shNC), two groups of nude mice received subcutaneous injections of shNC or shYTHDF1. The model with tumors, as well as the tumor volume curve, tumor photos and weights, are shown in **Figure 4A-D**. The growth rate, volume, and weight of the tumors in the shYTHDF1 group were significantly lower than those in the shNC group, according to the data. HE staining was used to evaluate the morphological characteristics of the xenograft tissues, and the results revealed that the morphology and structure of the cells were similar to those of the LUAD tissues **(Figure 4E)**. Immunohistochemical staining of Ki67 revealed that the proliferation ability of cells was significantly inhibited in the shYTHDF1 group compared with the shNC group **(Figure 4F, G)**. These findings indicate that YTHDF1 knockout can inhibit the growth of LUAD *in vivo*.

**Figure 4.**
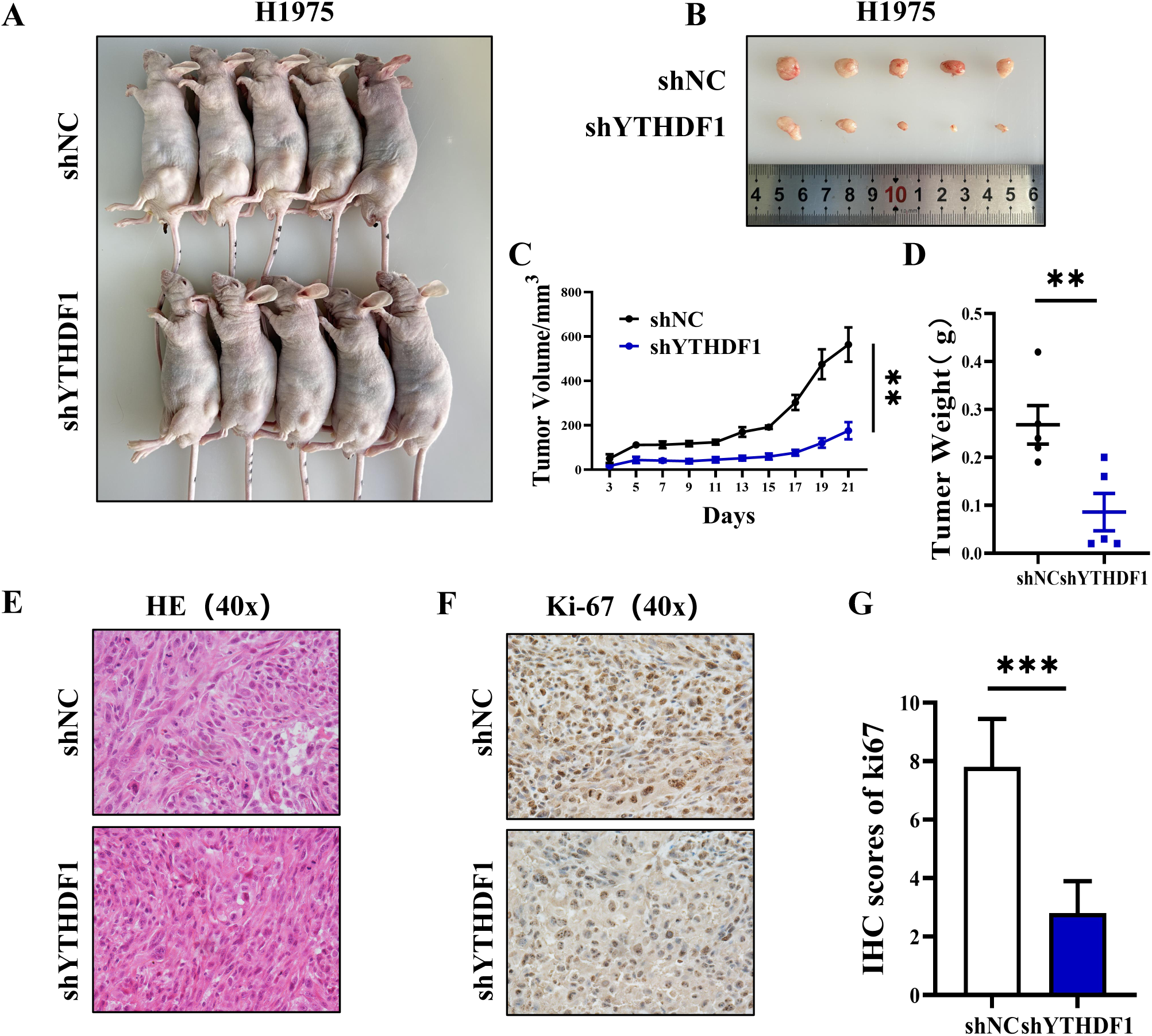
YTHDF1 regulates the growth of LUAD *in vivo*. **(A)** Nude mouse tumor-bearing model. **(C)** Tumour growth curve. **(B, D)** Images (B) and weights **(D)** of xenograft tumors were analysed. **(E)** Representative images of tumor tissue sections stained with HE. Scale bar, 50 μm. **(F)** Representative image of Ki-67 IHC staining of tumor tissue sections. Scale bar, 50 μm. The data are shown as the means ± SDs. \*\**P* <0.01, \*\*\**P* <0.001.

### Identification of candidate target genes with m6A modification of YTHDF1 in LUAD

To explore the potential mechanism of YTHDF1 in LUAD and screen for downstream target genes with m6A modification sites that interact with YTHDF1, we first performed RNA-seq and m6A-seq on H1975 cells with YTHDF1 knockdown and control cells. The RNA-seq results revealed a total of 210 significantly differentially expressed genes (*P*<0.05) between the YTHDF1-knockdown group and the control group, with 167 downregulated genes and 43 upregulated genes **(Figure 5A)**. The m6A-seq results revealed that the number of m6A peaks in the control and YTHDF1-knockdown H1975 cells was 8557 and 9104, respectively **(Figure 5B)**. Using the Homer motif discovery tool, we found that the “GGAC” consensus sequence was the predominant motif enriched in the m6A peaks **(Figure 5C)**, and these peaks were located within protein-coding transcripts enriched in CDSs and 3’UTRs, particularly near stop codons **(Figure 5D)**. Previous studies have reported that YTHDF1 does not affect the RNA abundance of its targets but rather regulates protein synthesis by interacting with m6A-methylated mRNAs. To screen for target genes with m6A methylations that can directly bind to YTHDF1, we further sequenced RNA from YTHDF1 immunopurified complexes (RIP-seq) in YTHDF1-knockdown and control H1975 cells to identify YTHDF1-bound mRNAs. The RIP-seq results revealed that YTHDF1-bound m6A peaks were also enriched mainly in CDSs and 3’UTRs, especially near stop codons **(Figure 5E)**, with the number of m6A peaks in the control and YTHDF1-knockdown H1975 cells being 19764 and 10339, respectively **(Figure 5F)**. Previous research has demonstrated that YTHDF1 knockdown does not significantly affect the m6A levels of its RNA; therefore, we next screened 7845 genes with significant m6A methylations in both groups of samples in the m6A-seq and 6051 genes that no longer bound to YTHDF1 after being knocked down via RIP-seq **(Figure 5G)**. Analysis of overlapping genes from RNA-seq, m6A-seq, and RIP-seq data revealed 1757 genes that bound to YTHDF1 and did not show significant changes in transcription levels after YTHDF1 knockdown, as indicated by m6A **(Figure 5H)**. GO functional enrichment analysis was re-performed on these 1757 genes **(Figure 5I)**, and 182 genes related to mRNA metabolism regulation and translation process regulation were identified (Q<0.05). We further identified 13 candidate target genes that are highly expressed in LUAD tissues according to the TCGA database; these genes are expressed mainly in the cytoplasm and are significantly negatively correlated with the overall survival rate of LUAD patients. Finally, via Integrative Genomics Viewer (IGV) software, we detected significant m6A peaks and YTHDF1 binding enrichment in 3 candidate genes (FBXO32, ERBB2, and EEF1G) **(Figure 5J)**. RIP-PCR was performed in H1975 cells, and the results revealed that YTHDF1 could bind to the transcripts of these 3 candidate genes, with EEF1G having the highest relative enrichment for YTHDF1 **(Figure 5K, L)**. In summary, EEF1G is the most likely candidate target gene that can directly bind to YTHDF1 and has m6A methylation. We further validated the expression level of EEF1G in LUAD tissues and cells and investigated the interaction between EEF1G and YTHDF1.

**Figure 5.**
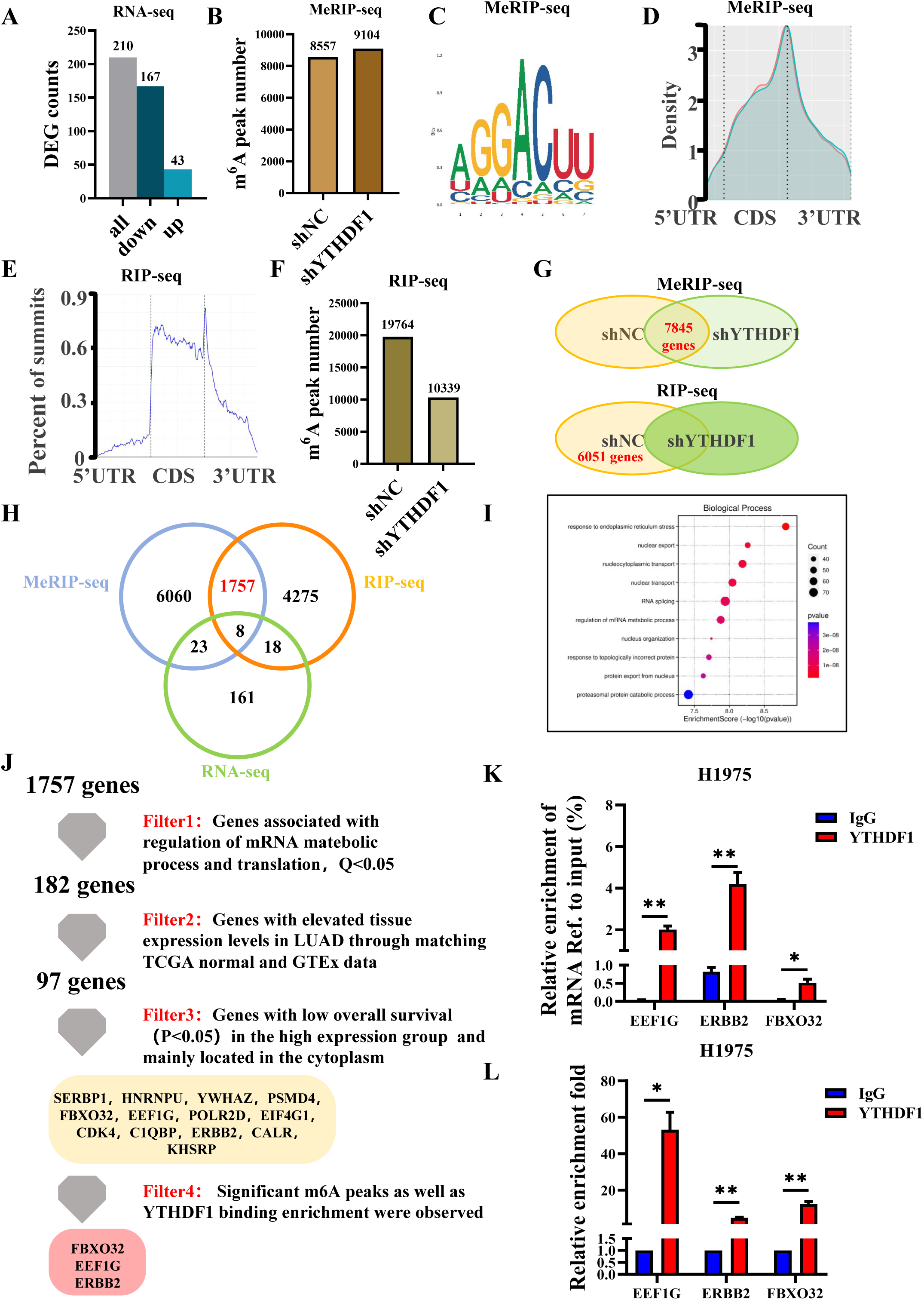
Identification of candidate target genes with m6A modification of YTHDF1 in LUAD. **(A)** Number of DEGs identified via RNA-seq. **(B-D)** The numbers of m6A peaks (B) and m6A motifs (C) and the density distributions of the m6A peaks in the 5’UTRs, start codons, CDSs, stop codons, and 3’UTRs (D) detected via MeRIP-seq. **(E, F)** Proportion of the m6A peak distribution in the 5’UTR, start codon, CDS, stop codon, 3’UTR (E), and number of differentially expressed genes (F) detected via RIP-seq. **(G)** The same genes were found in the MeRIP-seq siNC and siYTHDF1 groups, and the genes in the RIP-seq siNC group were different from those in the siYTHDF1 group. **(H)** Venn diagram illustrating the overlapping genes identified by m6A-seq, RIP-seq, and RNA-seq. **(I)** The top 10 biological processes in which 1757 genes are involved are shown by their Gene Ontology (GO) enrichment analysis. **(J)** Flow chart of the 3 selected candidate YTHDF1 target genes in H1975 cells. **(K, L)** RIP-PCR assays detecting the interactions between YTHDF1 and the mRNAs of 3 candidate genes in H1975 cells. IgG was used as an internal control.

### EEF1G is highly expressed in LUAD and directly interacts with YTHDF1

First, through the analysis of the online databases GEPIA2, GeneCards, and Kaplan‒Meier Plotter, we verified that EEF1G is highly expressed in LUAD tissues and is primarily distributed in the cytoplasm and that LUAD patients with high EEF1G expression have a lower overall survival rate **(Figure 6A‒C)**. Analysis with IGV software revealed significant m6A peaks and YTHDF1 binding enrichment for EEF1G **(Figure 6E)**. We further explored the potential binding sites of EEF1G with YTHDF1 through the online analysis website catRAPID and found that the YTHDF1 binding sites were consistent with the m6A modification sites **(Figure 6D)**. Next, we validated the mRNA and protein levels of EEF1G in normal lung bronchial epithelial BEAS-2B cells and three LUAD cell lines (H1975, H1299, and A549) via qRT‒PCR and Western blotting, respectively **(Figure 6F, G)**, and verified the protein expression level of EEF1G in LUAD tissues and adjacent normal tissues via IHC **(Figure 6H)**. The results revealed that EEF1G was highly expressed in LUAD cells and tissues (all *P*<0.05). In addition, we verified via Co-IP and Co-IF that YTHDF1 and EEF1G can coprecipitate in H975 cells and are located primarily in the cytoplasm of LUAD cells **(Figure 6I, J)**. Finally, to verify the regulatory role of YTHDF1 in the expression of EEF1G at the transcriptional and translational levels in LUAD, we detected the mRNA and protein expression levels of EEF1G in H1975 cells transfected with siNC/siYTHDF1 and vector/YTHDF1 through qRT‒PCR and Western blotting, respectively. After the expression level of YTHDF1 was downregulated or upregulated in H1975 cells, the mRNA level of EEF1G did not change significantly; however, the protein level of EEF1G was significantly downregulated or upregulated accordingly **(Figure 6K, L, M, N)**. These results suggest that YTHDF1 can interact with EEF1G and regulate its protein level at the translational level. To this end, we determined that EEF1G can serve as a m6A modification target of YTHDF1, and further studies should be conducted to explore the specific mechanisms by which YTHDF1 combines with EEF1G in the occurrence and development of LUAD.

**Figure 6.**
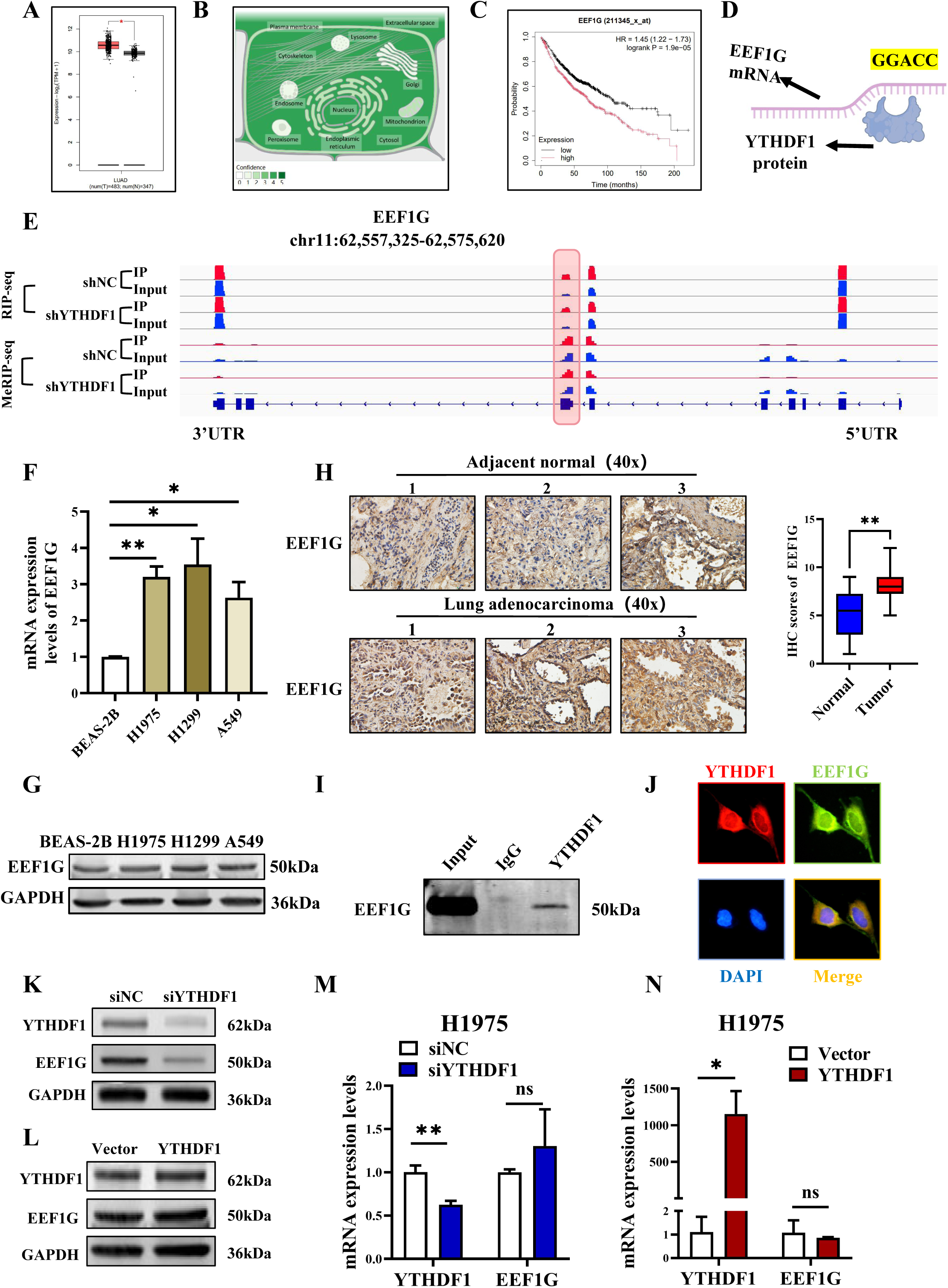
EEF1G is highly expressed in LUAD and directly interacts with YTHDF1. **(A)** Expression of EEF1G in LUAD samples determined by matching TCGA normal and GTEx data. **(B)** Subcellular localization of EEF1G in GeneCards. **(C)** EEF1G expression level and overall survival of LUAD patients: Kaplan‒Meier analysis. **(D)** The binding site of the YTHDF1 protein and EEF1G mRNA was predicted via the catRAPID website. **(E)** The distribution of m6A peaks and YTHDF1-binding peaks in EEF1G displayed by IGV tracks according to m6A-seq and YTHDF1 RIP-seq of H1975 cells. **(F)** RT‒qPCR was used to detect the relative mRNA levels of EEF1G in BEAS-2B, H1975, H1299, and A549 cells. **(G)** Western blot analysis of the protein levels of EEF1G in BEAS-2B, H1975, H1299, and A549 cells. **(H)** Representative IHC images of EEF1G protein expression in LUAD tissues and adjacent normal tissues. Scale bar, 50 μm. Quantitative analysis of EEF1G expression in LUAD tissues and adjacent normal tissues was performed in a double-blinded manner. **(I)** Co-IP followed by Western blot analyses confirmed the binding between YTHDF1 and EEF1G in H1975 cells. **(J)** YTHDF1 and EEF1G were colocalized mainly in the cytoplasm of H1975 cells, as demonstrated by fluorescence microscopy. The data are shown as the means ± SDs. \**P* <0.05, \*\**P* <0.01, \*\*\**P* <0.001. **(K, L)** Western blot analysis of the protein levels of EEF1G in H1975 cells upon YTHDF1 knockdown (K) or overexpression (L). **(M, N)** Relative mRNA levels of EEF1G in H1975 cells upon YTHDF1 knockdown (M) or overexpression (N).

### The oncogenic role of YTHDF1 depends on EEF1G in LUAD cells

To further study the role of EEF1G in LUAD and to evaluate whether the m6A reader YTHDF1 mediates the progression of LUAD by targeting EEF1G, we transfected siEEF1G into A549 cells (with the highest expression of EEF1G) where YTHDF1 was overexpressed and transfected a plasmid containing the EEF1G sequence into H1975 cells (with the lowest expression of EEF1G) where YTHDF1 was knocked down. We did this to determine whether the knockdown or overexpression of EEF1G could reverse the changes caused by YTHDF1 overexpression or silencing. The qRT‒PCR results revealed that the knockdown of EEF1G in A549 cells significantly reversed the increase in the mRNA level caused by YTHDF1, and the overexpression of EEF1G in H1975 cells significantly reversed the decrease in the mRNA level caused by YTHDF1 **(Figure S1A, B)**. The western blot results revealed that the knockdown of EEF1G in A549 cells significantly reversed the increase in protein levels caused by YTHDF1, and the overexpression of EEF1G in H1975 cells significantly reversed the decrease in protein levels caused by YTHDF1 **(Figure S1C)**. These results suggest that EEF1G can affect the expression of YTHDF1 in LUAD cells. We subsequently investigated whether EEF1G affects the impact of YTHDF1 on the biological functions of LUAD cells. After EEF1G was knocked down in A549 cells, the proliferative ability of LUAD cells was inhibited, as measured by the CCK-8 and colony formation assays **(Figure 7A, C)**; the migratory ability was inhibited, as shown by the wound healing and transwell migration assays **(Figure 7E, G)**; and the invasive ability was also inhibited, as shown by the transwell invasion assay **(Figure 7I)**. After overexpressing EEF1G in H1975 cells, the proliferative ability of LUAD cells was restored, as measured by the CCK-8 assay and plate colony formation assay **(Figure 7B, D)**, the migratory ability was restored, as shown by the wound healing and Transwell migration assays **(Figure 7F, H)**, and the invasive ability was restored, as shown by the Transwell invasion assay **(Figure 7J)**. These results indicate that EEF1G is a key downstream target of YTHDF1 in promoting the progression of LUAD and that the oncogenic role of YTHDF1 in LUAD cells depends on EEF1G. In summary, we believe that the m6A reader YTHDF1 can play a carcinogenic role in the pathogenesis of LUAD by recognizing the m6A site on EEF1G mRNA, promoting the translation of EEF1G, and increasing its protein expression.

**Figure 7.**
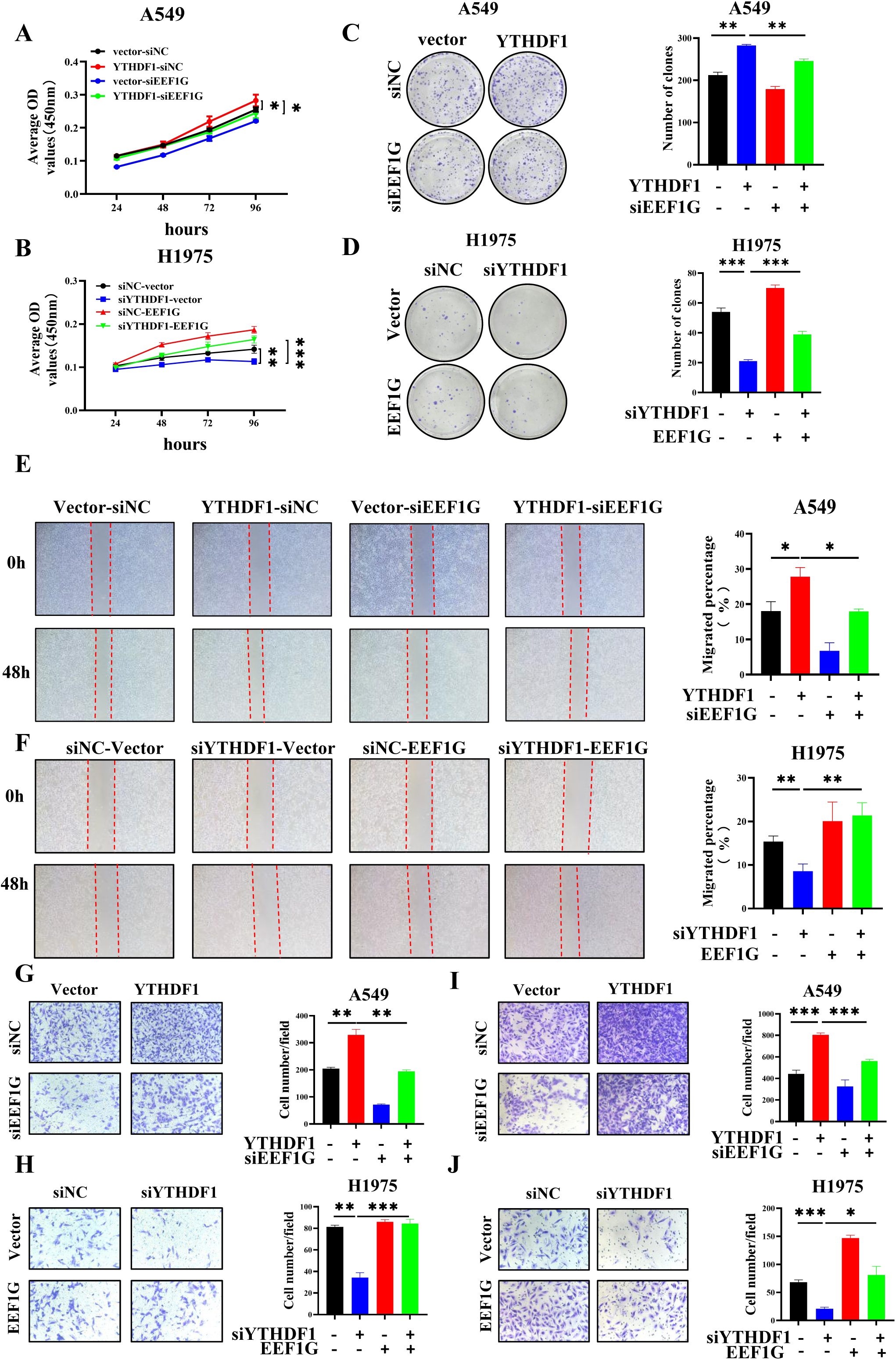
The oncogenic role of YTHDF1 depends on EEF1G in LUAD cells. **(A-J)** Effects of EEF1G on YTHDF1 in A549 and H1975 cells determined via the CCK-8 assay, colony formation assay, wound healing assay, Scale bar, 200 μm, Transwell migration assay, and Transwell invasion assay. Scale bar, 100 μm. EEF1G knockdown in A549 cells significantly reduced the effects of YTHDF1 on promoting cell viability (A), cell clonogenicity (C), cell migration ability (E, G), and cell invasiveness (I). Overexpression of EEF1G in H1975 cells significantly reduced the suppressive effects of YTHDF1 on cell viability (B), cell clonogenicity (D), cell migration ability (F, H), and cell invasiveness (J). The data are shown as the means ± SDs. ns, not significant; \**P* <0.05, \*\**P* <0.01, \*\*\**P* <0.001.

## Discussion

Dysregulation of m6A modification-related enzymes has been demonstrated to play crucial roles in cancer initiation, progression, metastasis, metabolism, drug resistance, and immune evasion, as well as in cancer stem cell self-renewal and the tumor microenvironment, highlighting the potential therapeutic value of targeting the m6A modification machinery for cancer treatment^6^. Recent studies have shown that METTL3-mediated m6A modification of SLC7A11 is regulated by the recruitment of YTHDF1, affecting the stability and translation of SLC7A11 mRNA and promoting LUAD cell proliferation while inhibiting cellular ferroptosis^15^. Another study revealed that the Wnt/β-catenin signalling pathway, by downregulating FTO expression, enhances the m6A modification of MYC mRNA, ultimately promoting glycolysis and tumorigenesis in lung cancer cells^16^. Furthermore, research has elucidated the mechanism by which YTHDF2 promotes LUAD cell proliferation and metastasis by regulating AXIN1 and the Wnt/β-catenin signalling pathway^17^. SUMOylation of YTHDF2 can increase its binding affinity with m6A-modified RNA, promoting the degradation of m6A-modified RNA and ultimately accelerating lung cancer progression^18^. Studies have confirmed that YTHDF1 is highly expressed in patients with KRAS/tp53 mutations and is associated with poor prognosis in LUAD patients through its promotion of cyclin B1 mRNA translation^19^. Furthermore, DUSP5, which is regulated by YTHDF1-mediated m6A modification, promotes epithelial‒mesenchymal transition and EGFR‒TKI resistance in LUAD through the TGF-β/Smad signalling pathway^20^. Despite the established oncogenic role of YTHDF1, the molecular mechanisms underlying how YTHDF1-mediated m6A modification participates in translational regulation through translation elongation factors, ultimately leading to the development of LUAD, remain unclear. Clarifying the specific molecular mechanisms of LUAD initiation and progression and identifying reliable therapeutic targets for LUAD are urgent tasks.

Reportedly, YTHDF1, a m6A-binding protein, is overexpressed in various tumors and is associated with poor prognosis. YTHDF1 exerts its oncogenic effect through different mechanisms, such as promoting translation or regulating mRNA stability by binding to m6A-modified mRNAs^10^. For example, the overexpression of YTHDF1 in hepatocellular carcinoma tissues is correlated with poor prognosis^21^; in breast cancer tissues, high expression levels of YTHDF1 are positively correlated with tumor size, lymph node involvement, and distant metastasis^22^. In ovarian cancer, the overexpression of YTHDF1 is associated with poor prognosis, and it promotes tumor growth and metastasis by enhancing the translation of EIF3C in a m6A-dependent manner^23^. In head and neck squamous cell carcinomas, YTHDF1 regulates iron metabolism during tumor progression both *in vivo* and *in vitro*, interacting with the 3’UTR and 5’UTR of TRFC mRNA to positively regulate the translation of TFRC mRNA in a m6A-dependent manner^24^. YTHDF1 is highly expressed in prostate cancer (PCa) tissues and cells, and high levels of YTHDF1 are also associated with a relatively poor prognosis in PCa patients. RNA sequencing and functional experiments have shown that YTHDF1 promotes the progression of PCa by increasing the translation of TRIM44 mRNA^25^. As an oncogene, YTHDF1 promotes the progression and invasion of breast cancer cells by inducing glycolysis. YTHDF can promote tumor glycolysis by increasing PKM2 levels, ultimately promoting the tumor growth and invasion of breast cancer cells^26^.

In this study, we found that YTHDF1 is highly expressed in LUAD tissues and cells compared with adjacent normal tissues and normal lung epithelial cells and that this high expression is positively correlated with tumor stage and negatively correlated with differentiation, which is consistent with previous reports. We also validated the results obtained from our analysis of the TCGA database and GEO database (GSE43458). Next, we explored the effects of overexpressing and knocking down YTHDF1 in LUAD cell lines on the biological functions of LUAD cells. We found that upregulating YTHDF1 significantly promoted the proliferation, migration, and invasion of LUAD cells and inhibited their apoptosis, and the opposite effects were observed when YTHDF1 was downregulated. Further *in vivo* experiments also highlighted the promoting effect of YTHDF1 on the proliferation of LUAD cells. These results collectively elucidate the oncogenic role of YTHDF1 in LUAD and its association with poor prognosis in LUAD patients, suggesting that YTHDF1 may represent a novel target for cancer therapy and prognosis. Therefore, m6A modification may play a crucial role in the progression of LUAD. Through a combination of RNA-seq, MeRIP-seq, and RIP-seq with bioinformatics analysis, we subsequently identified EEF1G as a direct target of YTHDF1. YTHDF1 primarily promotes the translation of EEF1G by binding to EEF1G mRNA in a m6A-dependent manner to exert its oncogenic effect.

Eukaryotic translation elongation factors (EEFs) play a central role in protein biosynthesis during the elongation step of translation. The elongation phase is carried out by the EEF1 and EEF2 complexes, with the subunits of the EEF1 complex, including EEF1A1, EEF1A2, EEF1B2, EEF1D, EEF1E1, and EEF1G, which are responsible for binding aminoacyl-tRNAs and transferring them to the A site of the ribosome^27^. In addition to their primary function within the EEF1 complex, recent studies have also attributed significant implications to these proteins in tumorigenesis mechanisms. Analysis of the human genome revealed the presence of nine pseudogenes for EEF1G, classified as processed pseudogenes, but these pseudogenes have been studied very marginally^28^. Most of them are predicted through genome sequence analysis and lack experimental evidence to support their existence. EEF1G is reportedly overexpressed in gastric cancer^29^, colorectal adenocarcinoma^30^, and pancreatic cancer^31^. Studies have also detected the overexpression of EEF1G in lung tumor tissues via the TCGA dataset, with increased transcript levels leading to poorer overall survival (OS) and first progression (FP) in patients with lung cancer^32^. Further histological subtype analysis revealed a significant correlation between increased EEF1G mRNA levels and poorer survival outcomes in patients with LUAD, which is consistent with our findings.

In this study, we found that EEF1G is highly expressed in LUAD tissues and cells and is significantly associated with poor patient prognosis, suggesting that EEF1G may act as an oncogene in the development of LUAD and has the potential to be a novel biomarker for predicting LUAD. Furthermore, the dysregulation of EEF1 genes and their expression profiles can affect epigenetic mechanisms in cancer, with at least one variant in the EEF1 gene affecting the levels of histone methyltransferases and acetyltransferases^33^. In this study, we elucidated a novel regulatory mechanism of EEF1G, which is controlled by YTHDF1 in a m6A-dependent manner. Through Co-IP and Co-IF experiments, we discovered a direct interaction between YTHDF1 and EEF1G in LUAD cells, which has not been reported in existing studies. Furthermore, bioinformatics analysis predicted the presence of a binding site between the YTHDF1 protein and EEF1G mRNA that aligns with the m6A modification site, further supporting our conclusions regarding their interaction. Consistent with previous studies, through in-depth exploration of the regulatory mechanism of YTHDF1 on EEF1G in LUAD, we confirmed that YTHDF1 controls the expression of EEF1G at the translational level without affecting its transcriptional level. Rescue experiments demonstrated that overexpression of EEF1G is sufficient to rescue the inhibitory effect of YTHDF1 gene downregulation on LUAD cells, whereas EEF1G knockdown inhibits the promoting effect of YTHDF1 gene upregulation on LUAD cells. These findings indicate that EEF1G is a reliable downstream target of YTHDF1 in LUAD and that YTHDF1 exerts its oncogenic effect in LUAD by promoting the translation of EEF1G mRNA through m6A modification.

In this study, we propose for the first time that EEF1G serves as a direct target of YTHDF1, with YTHDF1 promoting the translation of EEF1G mRNA in a m6A-dependent manner. However, our research has several limitations. At the molecular level, we have studied only the downstream targets of YTHDF1, and further investigations are needed to determine the specific pathways through which YTHDF1 regulates EEF1G expression. Additionally, we have not yet validated *in vivo* whether YTHDF1 could be a therapeutic target for LUAD. In future research, we aim to address these issues, further validate the specific molecular mechanisms by which YTHDF1 regulates EEF1G to promote the development of LUAD, and identify reliable therapeutic targets for the treatment of LUAD.

## Conclusion

In summary, our study suggested that YTHDF1, a m6A “reader,” enhances LUAD cell proliferation, migration, and invasion and inhibits apoptosis. EEF1G is a direct target of YTHDF1, which is regulated by YTHDF1 in a m6A-dependent manner and plays a significant role in LUAD. YTHDF1 may serve as a potential therapeutic target for LUAD.

## Author Contributions

LHW, QHS and XYW designed and performed the experiments; HJY, QW and MZ collected and validated the clinical sample data; JLM and LW extracted, analysed and interpreted the data from the GEO and TCGA databases. LHW, QHS and XYW wrote the manuscript. JW, TL and WFY made substantial contributions to the conception of the work and substantively revised it. JW, TL and WFY contributed to the study supervision. All the authors read and approved the final manuscript.

## Availability Statement

The data underlying this article will be shared upon reasonable request to the corresponding author.

## Supporting information

Additional supporting information may be found online in the Supporting Information section at the end of the article.

## Supplementary Figure

**Supplementary Figure 1**

**(A, B)** The expression of YTHDF1 mRNA in A549 (A) and H1975 (B) cells was measured by qRT‒PCR after cotransfection of the corresponding siRNA and overexpression plasmid. **(C)** The expression of the YTHDF1 protein in A549 and H1975 cells was detected by Western blotting after cotransfection with the corresponding siRNA and overexpression plasmid.

## Supplementary Table

**Supplementary Table 1** Primer sequences for YTHDF1, EEF1G, ERBB2, FBXO32, and β-actin.

