## Supplementary figure 1 for "YTHDF1 Facilitates Lung Adenocarcinoma Progression via Promotion of EEF1G Translation in a m6A-Dependent Manner"

### Slide 1
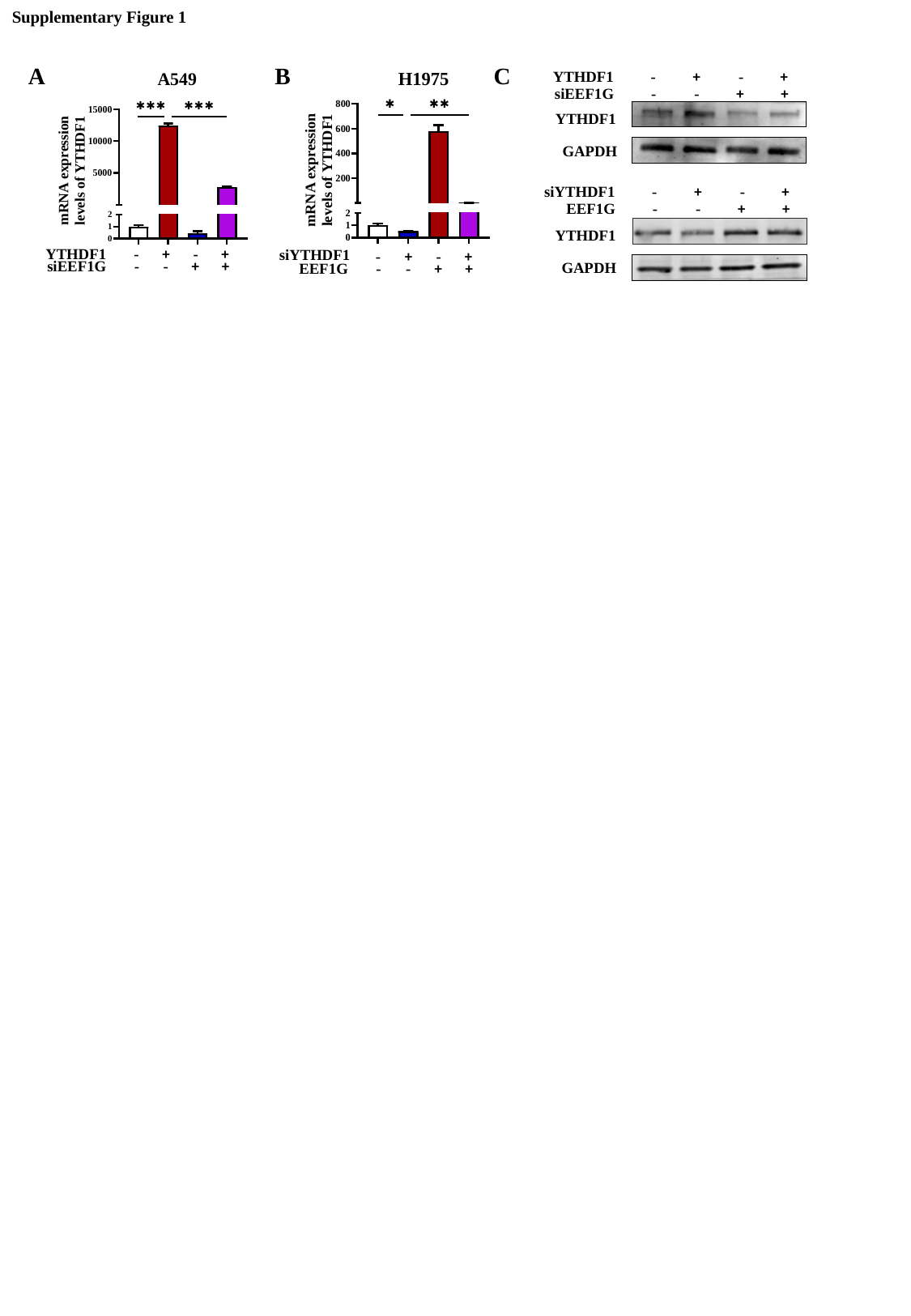

Supplementary Figure 1
A
B
C
siYTHDF1
-
+
-
+
-
-
+
+
EEF1G
YTHDF1
-
+
-
+
-
-
+
+
siEEF1G
-
+
-
+
YTHDF1
-
-
+
+
siEEF1G
YTHDF1
GAPDH
siYTHDF1
-
+
-
+
-
-
+
+
EEF1G
YTHDF1
GAPDH
