## Supplementary table 1 for "YTHDF1 Facilitates Lung Adenocarcinoma Progression via Promotion of EEF1G Translation in a m6A-Dependent Manner"

### Slide 1
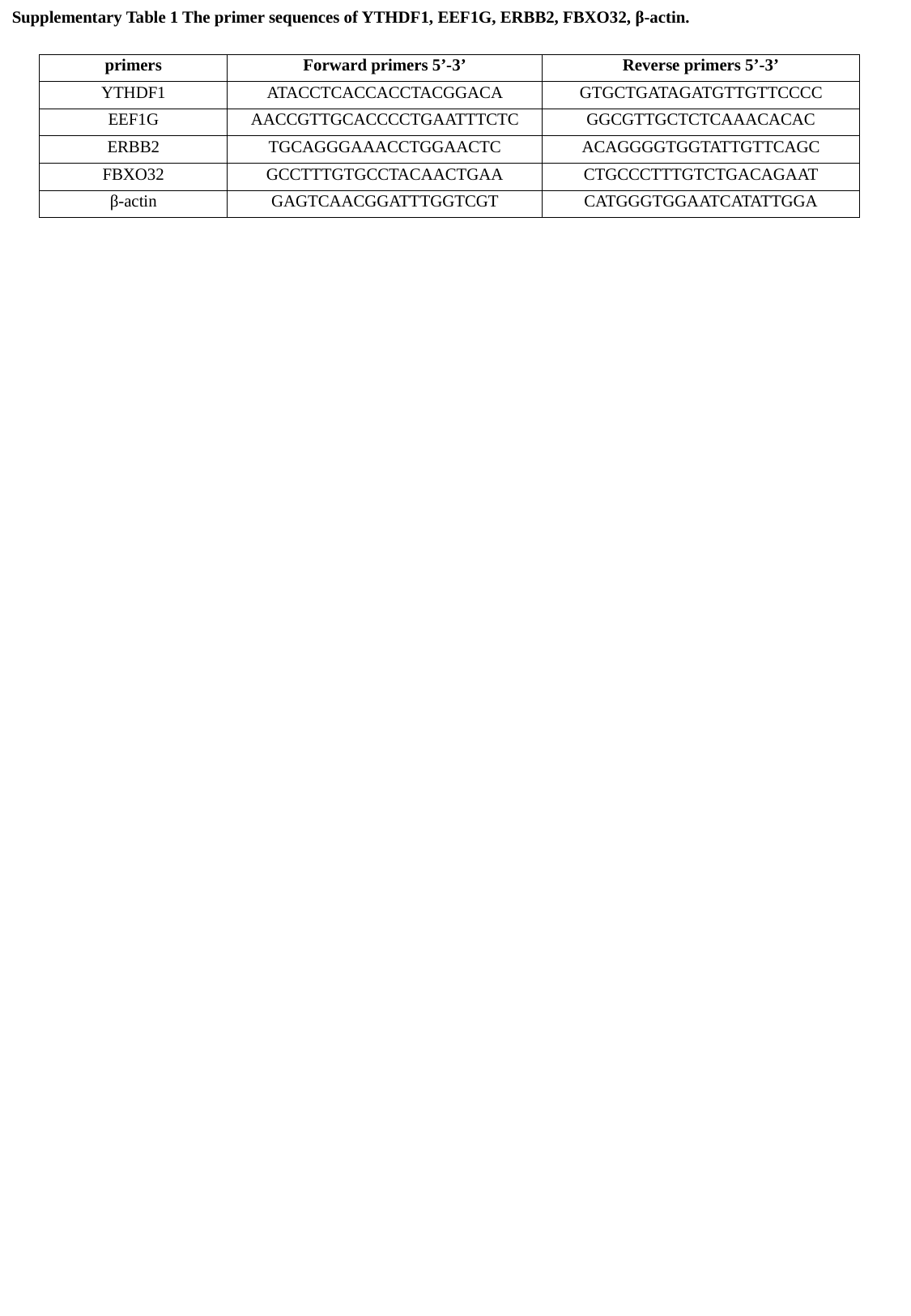

Supplementary Table 1 The primer sequences of YTHDF1, EEF1G, ERBB2, FBXO32, β-actin.
| primers | Forward primers 5’-3’ | Reverse primers 5’-3’ |
| --- | --- | --- |
| YTHDF1 | ATACCTCACCACCTACGGACA | GTGCTGATAGATGTTGTTCCCC |
| EEF1G | AACCGTTGCACCCCTGAATTTCTC | GGCGTTGCTCTCAAACACAC |
| ERBB2 | TGCAGGGAAACCTGGAACTC | ACAGGGGTGGTATTGTTCAGC |
| FBXO32 | GCCTTTGTGCCTACAACTGAA | CTGCCCTTTGTCTGACAGAAT |
| β-actin | GAGTCAACGGATTTGGTCGT | CATGGGTGGAATCATATTGGA |
